# Mature rotavirus particles contain equivalent amounts of ^7me^GpppGcap and noncapped viral positive-sense RNAs

**DOI:** 10.1101/2022.06.03.494783

**Authors:** Joaquin Moreno-Contreras, Liliana Sánchez-Tacuba, Carlos F. Arias, Susana López

## Abstract

Viruses have evolved different strategies to overcome their recognition by the host innate immune system. Addition of cap at their 5’RNA ends is an efficient mechanism to ensure escape from detection by the innate immune system, but also to ensure the efficient synthesis of viral proteins. Rotavirus mRNAs contain a type 1 cap structure at their 5’end that is added by the viral capping enzyme VP3. This is a multifunctional protein with all the enzymatic activities necessary to add the cap, and also functions as an antagonist of the OAS-RNase L pathway. Here, the relative abundance of capped and noncapped viral RNAs during the replication cycle of rotavirus was determined. We found that both classes of rotaviral +RNAs are encapsidated, and they were present in a 1:1 ratio in the mature infectious particles. The capping of viral +RNAs is dynamic since different ratios of capped and noncapped RNAs were detected at different times post infection. Similarly, when the relative amount of capped and uncapped viral +RNAs produced in an *in vitro* transcription system was determined, we found that the proportion was very similar to that in the mature viral particles and in infected cells, suggesting that the capping efficiency of VP3 both, *in vivo* and *in vitro,* might be close to 50%. Unexpectedly, when the effect of simultaneously knocking down the expression of VP3 and RNase L on the cap status of viral +RNAs was evaluated, we found that even though at late times post infection there was an increased proportion of capped viral RNAs in infected cells, the viral particles isolated from this condition contained an equal ratio of capped and noncapped viral RNA, suggesting that there might be a selective packaging of capped-noncapped RNAs.

**SIGNIFICANCE:** Rotaviruses have a genome composed of eleven segments of double stranded RNA. Whether all 5’ ends of the positive sense genomic RNA contained in the mature viral particles are modified by a cap structure is unknown. In this work, using a direct quantitative assay we characterized the relative proportion of capped and noncapped viral RNA in rotavirus infected cells and in viral particles. We found that independently of the relative proportions of cap/noncapped RNA present in rotavirus infected cells, there is a similar proportion of these two kinds of 5’-modified positive sense RNAs in the viral particles.

## INTRODUCTION

A signature of cellular mRNAs is the presence of N7-methyl-guanosine triphosphate (cap) at their 5’end. The cap structure is important for different mRNA cellular processes, including: splicing, nuclear-cytoplasmic export, translation initiation, stability, and recognition of self from foreign RNAs (1, 2). The canonical RNA capping pathway takes place in the nucleus and occurs co-transcriptionally by a sequential enzymatic process of three enzymes; an RNA triphosphatase (RTPase) that removes the *γ*-phosphate from 5’triphosphate to generate a 5’diphosphate end, an RNA guanylyltransferase (GTase) that binds a guanine monophosphate (GMP) nucleotide from GTP to the 5’diphosphate RNA to form a 5’guanylylated-RNA, and a guanine-N7 methyltranferase (N7-MTase) that transfers a methyl group from S-adenosylmethionine (SAM) to the N7 position of the terminal guanine base, completing the synthesis of the type 0 cap structure(3). Additionally, the RNA 2’-O-ribose methyltransferase adds a methionine to the ribose 2’-hydroxyl (2’-O) of the first nucleotide forming the type 1 cap structure.

In the celĺs cytoplasm, noncapped RNAs are sensed as non-self by cellular cytoplasmic pattern recognition receptors such as RIG-1 or MDA5, triggering a signaling cascade that ends with the expression of interferon, inducing an antiviral state (1). To overcome the recognition of their transcripts by proteins of the innate immune system viruses have evolved different strategies to hide the 5’end of viral mRNAs using different strategies such as hijacking of the cellular capping machinery, acquiring cap structures from cellular RNAs by “cap snatching”, or by encoding their own capping machinery (4, 5).

Rotaviruses are a leading cause of severe gastroenteritis in young animals, including humans, causing an estimate of 258 millions episodes of diarrhea and 128,500 deaths of children younger than 5 years annually, mainly in developing countries around the world (6). These nonenveloped viruses belong to the *Reoviridae* family and are formed by three protein layers that surround the genome composed of eleven segments of double-stranded RNA (dsRNA). The structural proteins VP4 and VP7 form the outer layer, the middle layer is composed by VP6, and the inner-most layer is formed by VP2, and small amounts of the viral RNA-dependent RNA polymerase (RdRp) VP1, and the methyl-guanylyl transferase VP3 (7).

After entering the cell, the viral outer layer is released yielding a double-layered particle (DLP) that is transcriptionally active, and begins the synthesis of the plus-sense, messenger viral RNAs (+RNAs), these mRNAs direct the synthesis of all viral proteins, and are also used as templates for the synthesis of the negative strand RNAs (-RNAs) to form the genomic dsRNA. Once a critical amount of viral proteins is made, the formation of cytoplasmic, electrodense structures called viroplasms begin; in these non-membranous structures the replication of the viral genome and the initial steps of the morphogenesis of virions take place (8). The assembly of new transcriptionally active DLPs in viroplasms initiates a second round of transcription, increasing the amount of viral RNAs in the infected cell (9).

Similar to cellular mRNAs, rotavirus mRNAs contain a cap type 1 (^m7^GpppGm) at their 5’end, a structure that is added by the 98-kDa viral protein VP3, encoded in dsRNA segment 3 of the rotavirus genome (10). Recently, Kumar et al. described a detailed cryo-electron microscopy structure of this protein that showed that VP3 forms a stable tetramer with a modular organization of five domains, four of them involved in cap addition (11). While previous studies demonstrated that VP3 had RNA binding, guanylyl and methyl transferase activities (8, 10), it was not clear whether the helicase and RNA triphosphatase (RTPase) activities were also present in VP3, or in another viral protein. In their work, Kumar et al. demonstrated that these last two activities are located in the phosphodiesterase (PDE) domain of VP3 (11, 12). The PDE domain is also involved in the degradation of the 2’-5’oligoadenylates, thus blocking the activation of RNase L, and inhibiting the innate immune OAS/RNaseL pathway (13, 14).

Thus far, it has not been characterized during the replication cycle of rotavirus whether the viral +RNAs that are translated (mRNAs) contain a cap structure, while the +RNAs that serve as templates for the negative strand synthesis are uncapped, or if all +RNAs synthesized in infected cells contain a 5’cap structure. To address this question, we sought to quantitatively evaluate the relative abundance of capped and noncapped viral RNAs during rotavirus replication. Here, we report that both, capped and noncapped rotaviral +RNAs, are present in a similar proportion in mature infectious particles. We also show that capping of viral +RNAs is dynamic since different ratios of capped and noncapped RNAs are found at different times post infection. Likewise, when the relative amount of capped and noncapped viral +RNAs produced in an *in vitro* transcription system was characterized, their proportion was very similar to that found in infected cells, suggesting that the capping efficiency of VP3 *in vivo* and *in vitro* is close to 50%.

Unexpectedly, in cells where VP3 was silenced, the specific infectivity of the viral particles produced was higher than in control cells, although they contained a similar ratio of capped and noncapped RNAs; as expected, the viral RNA synthesized *in vitro* by the VP3-deficient particles was poorly capped. Finally, when the effect of co-silencing VP3 and RNase L on the cap status of viral +RNA was evaluated, we found that there was an increased proportion of capped viral RNAs at late times of infection, but the viral particles isolated from this condition contained an equal ratio of capped and noncapped RNA, suggesting that there might be a selective packaging of capped-noncapped viral RNAs.

## MATERIALS AND METHODS

### Cell culture and virus

The rhesus kidney epithelial cell line MA104 (ATCC) was grown in Dulbecco’s modified Eagle’s medium-reduced serum (DMEM-RS; Thermo Scientific HyClone, Logan, UT) supplemented with 5% heat-inactivated fetal bovine serum (FBS; Biowest, Kansas City, MO) at 37°C in a 5% CO_2_ atmosphere and was used for all experiments carried out in this work. Simian rotavirus strain RRV was originally obtained from H.B. Greenberg (Stanford University, Stanford, CA) and was propagated in MA104 cells as described previously (16). Prior to infection, the RRV-containing cell lysates were activated with 10 μg/ml of trypsin (Gibco Life Technologies, Carlsbad, CA) for 30 min at 37°C.

### Reagents and antibodies

Small interfering RNAs (siRNAs) were purchased from GE Healthcare Dharmacon (Lafayette, CO). The sequences of the siRNAs against rotavirus dsRNA segment 3 that encodes for VP3 have been previously reported (17). As an irrelevant control, the siGENOME nontargeting siRNA 4 from Dharmacon was used. The monoclonal antibody (MAb) to RNase L was obtained from Abcam (Cambridge, MA) and the MAb to VP3 was purchased from GeneScript. The rabbit anti-rotavirus polyclonal serum raised against purified triple-layered particles (TLPs), as well as the rabbit polyclonal sera to human vimentin (anti-Vim) were produced in our laboratory (18). Horseradish peroxidase-conjugated goat anti-rabbit polyclonal and anti-mouse antibodies were purchased from PerkinElmer Life Sciences (Boston, MA).

### Infectivity assay

MA104 cells were grown in 96-well plates to confluence, and infected with 2-fold serial dilutions of the virus cell lysates for 1 h at 37°C. After this time, the unbound virus was removed, and the cells were incubated in MEM without serum for 14 h at 37°C. Afterwards, the cells were fixed with 80% acetone in PBS for 30 min at room temperature and then washed twice with PBS. The fixed monolayers were then incubated with a rabbit anti-rotavirus polyclonal serum, followed by incubation with a secondary anti-rabbit antibody conjugated with horseradish peroxidase. Finally, the cells were washed twice with PBS and stained with a solution of 1 mg/ml of carbazole (3-amino-9-ethylcarbazole [AEC]) in sodium acetate buffer (50 mM, pH 5) with 0.04% H_2_O_2_. The reaction was stopped by washing in tap water, and the infectious focus-forming units were counted in a Nikon TMS inverted phase-contrast microscope with a 20X objective (19).

### siRNAs transfection

Transfection and co-transfection of siRNAs into MA104 cells was performed using Oligofectamine reagent (Invitrogen, Carlsbad, CA) in 48- or 6-well plates using a reverse transfection method as described previously (20).

### Immunoblot assays

Cells were lysed in Laemmli sample buffer and denatured by boiling for 5 min. Proteins in cell lysates were separated by 10% SDS-PAGE and transferred to Immobilon NC membranes (Millipore Merck KGaA, Darmstadt, Germany). The membranes were blocked by incubation with 5% nonfat dry milk in phosphate-buffered saline (PBS) overnight at 4°C and then incubated with primary antibodies diluted in PBS containing 0.1% nonfat dry milk. The membranes were then incubated with the corresponding secondary antibodies conjugated to horseradish peroxidase. The peroxidase activity was developed using the Western Lightning Chemiluminescence Reagent Plus (PerkinElmer Life Sciences, Boston, MA) according to the manufacturer’s instructions. The blots were also probed with an anti-vimentin antibody (anti-Vim), which was used as a loading control.

### TLP and DLP purification

MA104 cells grown in 150-cm^2^ flasks were infected with RRV at a multiplicity of infection (MOI) of 3 and, at 12 h post infection (hpi), cells were lysed by three cycles of freeze-thaw, and the viral particles were concentrated by ultracentrifugation for 1h at 30,000 rpm at 4°C in an SW40 rotor (Beckman, Fullerton, CA). Viral pellets were resuspended in TNC buffer (10 mM Tris-HCl [pH 7.5], 140 mM NaCl, 10 mM CaCl2), sonicated three times for 20 s and extracted with trichloromonofluoromethane as previously described (19). To purify viral TLPs, CsCl was added to the aqueous phase obtained from the trichloromonofluoromethane extraction to obtain a final density of 1.36 g/cm^3^, and the mixture was centrifuged for 18 h at 36,000 rpm in an SW55 Ti rotor (Beckman, Fullerton, CA) at 4°C. The opalescent bands corresponding to TLPs or DLPs were collected by puncture and stored at 4°C. Before use, viral particles were desalted in a Sephadex G-25 spin column and resuspended in TNC buffer. Purified TLPs were treated with 10 μg of RNase A for 15 min at 37°C prior to dsRNA extraction with TRIzol (Invitrogen, Carlsbad, CA) reagent as indicated by manufacturer.

To obtain viral particles from infected cells in which VP3 and RNase L were either silenced individually or co-silenced, MA104 cells grown in 6-wells plates were transfected with the indicated siRNA (13). At 72 h post transfection (hpt), cells were infected with RRV at an MOI of 3 and were harvested 12 hpi. Viral lysates were freeze-thawed three times, sonicated and then extracted with trichloromonofluoromethane as described above. The virus suspension was pelleted through a 40% sucrose cushion by centrifugation for 2 h at 40,000 rpm in a SW55 Ti rotor (Beckman, Fullerton, CA) at 4°C. The viral pellet was resuspended in TNC buffer (for total viral particles) or in TNE buffer (10 mM Tris-HCl [pH 7.5], 140 mM NaCl, 5 mM EGTA) to remove the outer layer proteins of the particles to obtain DLPs.

### *In vitro* transcription

*In vitro* transcription (IVT) reactions were carried out as follows: 2.4 μg of purified DLPs were mixed in a reaction mixture containing 8 mM ATP, 2.5 mM GTP, 2.5 mM CTP, 2.5 mM UTP (Ambion, Carlsban, CA), 0.5 mM S-adenosylmethionine (SAM, NEB, Ipswich, MA), 0.5 U/μl RNasin (Thermo Scientific, Waltham, MA), Transcription Buffer (140 mM Tris-HCl pH 8, 200 mM sodium acetate, and 40 mM magnesium acetate) and a 4% suspension of bentonite. Reaction mixtures were incubated for 6 h at 42°C. The reaction mixture was centrifuged in an Eppendorf centrifuge for 1 min at 14,000 rpm to remove the bentonite and the RNA present in the supernatant was immediately isolated by extraction with phenol:chloroform and precipitated with 4M LiCl overnight at 4 °C. Then, RNA was sedimented by centrifugation, washed with 80% ethanol, dried briefly and resuspended in RNase-free water. RNA concentration was determined by absorbance on a NanoDrop spectrophotometer and stored in aliquots at −70 °C.

### RNA preparation

MA104 cells were grown in 6-well plates to confluence and infected with rotavirus strain RRV at an MOI of 10. At different hpi total RNA was isolated with TRIzol as indicated by the manufacturer. Viral RNA from IVT, genomic dsRNA from viral particles, and RNA from infected cells, were obtained as described above. To characterize the 5’ends of viral RNAs, the following amounts of input RNA were employed: 250 ng of purified RNAs obtained from either genomic dsRNA extracted from viral particles or RNA from IVT reactions; and 1 μg of total RNA obtained from infected cells.

### Characterization of the 5’end of viral RNA

To characterize the 5’end of the plus sense rotavirus RNAs (+RNAs), we carried out an enzymatic process followed by RT-qPCR as previously described (15), with the following modifications (Fig. 1A): The input RNAs were treated with Antarctic Phosphatase (NEB, Ipswich, MA) for 1 h at 37°C to remove any 5’end phosphates. Then, the reaction mixture was heat-inactivated and split into two equal aliquots to assay the noncapped and ^7me^GpppG capped RNAs in parallel. For the noncapped assay the RNA in the reaction was treated with T4 PNK (NEB, Ipswich, MA) to produce a pool of 5’ monophosphate RNAs, and ^7me^GpppG capped RNAs. The other aliquot was treated with RppH (NEB, Ipswich, MA) to remove the ^7me^GpppG from capped RNAs, producing a pool of 5’ non-phosphorylated and decapped 5’-monophosphate RNAs. Both reactions were incubated during 1 h at 37°C, and afterwards the enzymes were heat-inactivated. Then, both reactions were used in parallel as substrates for the ligation with an RNA oligonucleotide (named linker RNA, 5’-GCUGAUGGCGAUGAAUGAACACUGCGUUUGCUGGCUUUGAUGAAA-3’), with the 5’ monophosphates available on the treated viral RNAs using T4 RNA ligase (NEB, Ipswich, MA) for 1 h at 37°C. After the ligation step, samples were extracted with phenol:chloroform, ethanol precipitated and resuspended in RNase free H2O.

**Figure 1.**
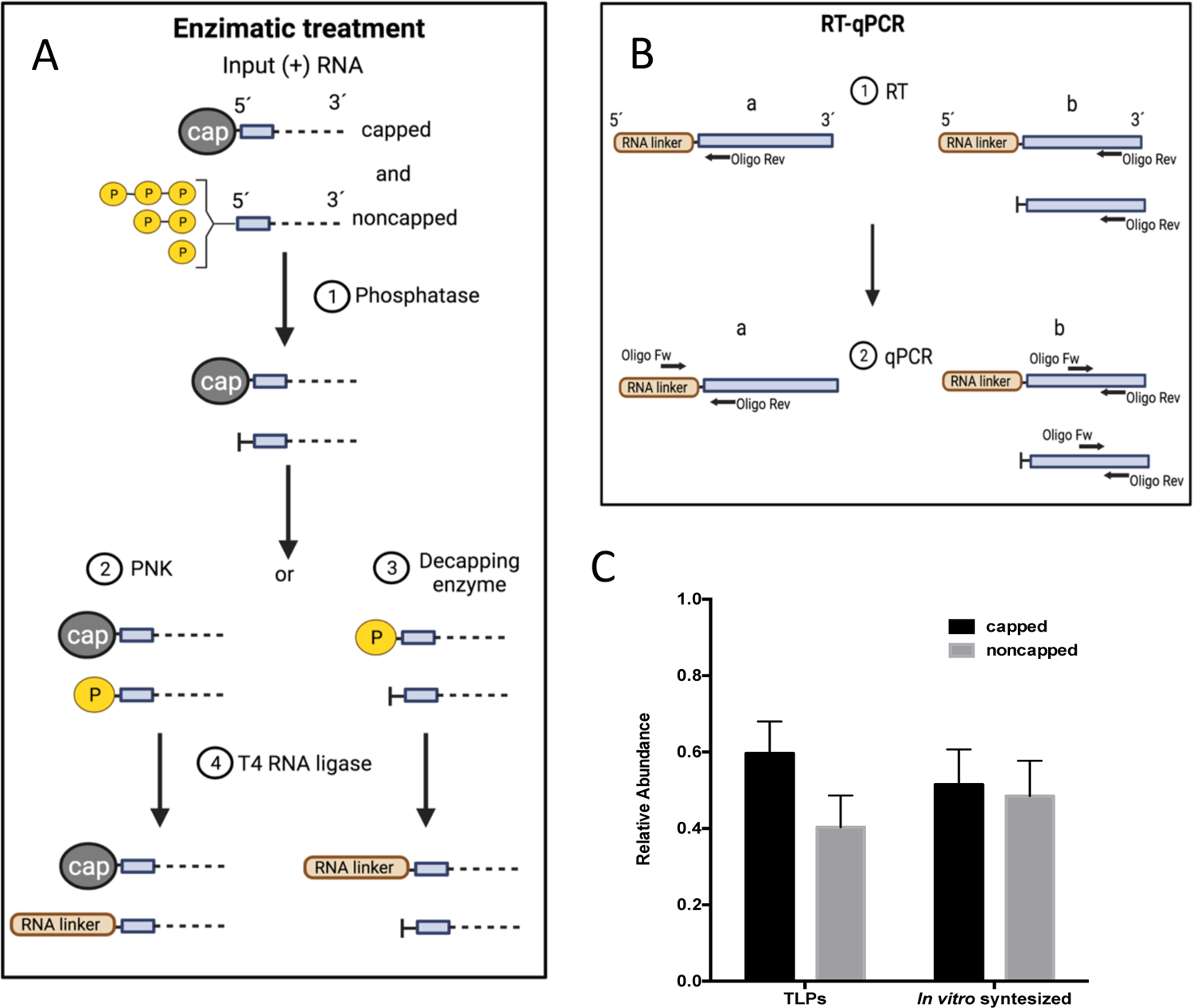
Quantitative determination of capped and noncapped viral RNA. Schematic representation of the method used for the quantitative capping determination on viral 5’ends. The input RNA consists of either genomic dsRNA, or viral ss+RNA. A) **Enzymatic treatment** (1) Capped or noncapped ss+RNAs were treated with Antarctic phosphatase, and then the reaction was split in two. (2) One half was treated with PNK, that adds a phosphate group to the 5’OH RNA. (3) The other half was treated with RppH; that cleaves the cap from RNA molecules leaving a 5’phosphate group. (4) An RNA linker was ligated to the free phosphate groups present at 5’end of RNA molecules using T4 RNA ligase. B) **RT-qPCR reactions.** Two RT-qPCR reactions were performed. Specific oligonucleotides were employed to perform the reverse transcription (RT) reaction to detect the amplicon produced by the ligation of the RNA linker with the viral segment 10 (a), or to detect the total amount of viral gene 10 in the reaction (b). All reactions were performed as detailed in material and methods section. C) Quantitative determination of capped and noncapped RNAs in mature viral particles (TLPs), and in *in vitro* synthetized viral RNAs. The arithmetic means +/- standard deviation from three independent biological replicates are shown.

### Reverse Transcriptase Reaction

The reverse transcription (RT) quantitative-PCR (RT-qPCR) was performed in two steps as described by Ayala-Breton *et al* (9). After purification, the RNAs were used as substrates for an RT reaction using specific oligonucleotides complementary to the positive strand of RNA segment 10 (Fig. 1 B, step 1). cDNA was reverse transcribed using M-MLV reverse transcriptase (Invitrogen, Waltham, MA) with the following oligonucleotides; to detect the linker-containing amplicon a reverse oligonucleotide (5’-CGAGGCCTTATGTAATGTAAATAG-3’) complementary to the 5’-end of viral segment 10 at positions 168 to 191 was employed (Fig.1B step 1 [a]); meanwhile, to quantify the total amount of viral segment 10 present in the reaction, the following inner reverse oligonucleotide (5’-GAGCAATCTTCATGGTTGGAA-3’) that binds at the positions 214 nt to 193 nt of viral segment 10 was used (Fig.1B step 1 [b]).

### Real time RT-qPCR analysis

After cDNA synthesis, the following oligonucleotides were used to detect the linker-containing amplicon. The forward oligonucleotide that binds on the RNA linker (5’-CGATGAATGAACACTGCGTTTG-3’) and the reverse oligonucleotide (5’-CGAGGCCTTATGTAATGTAAATAG-3’) that binds at the 5’end of gene 10 were used (Fig.1 B step 2 [a]). In parallel, primers used for the amplification of an inner region of rotavirus RNA segment 10, described previously, were used as an internal control of each amplification (9) (Fig.1 B step 2 [b]); the forward primer (5’-TCCTGGAATGGCGTATTTTC-3’) and the reverse primer (5’-GAGCAATCTTCATGGTTGGAA-3’) were used in the qPCR reaction. Samples were analyzed in an ABI 7000 sequence detector system (Applied Biosystems). To perform quantitative analysis of the data, the results were normalized to the levels of rotavirus segment 10 detected in each RNA sample. The changes in gene expression were calculated by the 2^-ΔΔCT^ method, were CT is the threshold cycle, as previously described (15, 21). Finally, the relative amounts of 5’capped and noncapped viral RNA was determined by comparing the relative quantities of the linker containing amplicon with the total amount of viral +RNA present, using the ΔΔCT method to obtain the ratios of 5’capped and noncapped RNAs depending on the sample treated, as previously described (15).

### Determination of the viral gene 10 copy number

To determine the number of viral copies in a sample, a standard curve was generated using a 10-fold serial dilution of an *in vitro* T7 transcribed RNA that contains the sequence of rotavirus RRV segment 10. Briefly, the logarithm of the concentration of each dilution was plotted against the C_T_ and the viral copy number from unknown samples was determined by extrapolating the CT value onto the corresponding standard curve.

### Statistical analysis

The data reported in this study represent the means of a minimum of three independent biological replicates. Error bars indicate the +/- standard deviation of the means. Statistical analysis was performed using GraphPad Prism 6.0 (GraphPad Software Inc) as described in Results.

## RESULTS

### The genomic RNA segments within the mature rotavirus particles contain capped and noncapped positive-sense RNA strands

The rotavirus genome, formed by eleven dsRNA segments is enclosed in viral particles, however, whether the positive strand of the genomic dsRNA contained in the mature viral capsid is capped or not, has not been characterized (22, 23). To quantitatively evaluate the proportion of the 5’cap present on rotaviral RNAs, we resorted to an enzymatic method for cap quantification that was previously described to determine the presence of cap in the RNA of Sindbis virus (SINV) (15). In this method, total RNA is treated with a phosphatase to remove 5’terminal phosphates. The sample is then divided into two equal aliquots; one is treated with polynucleotide kinase (PNK) that introduces a phosphate group in the 5’end of uncapped RNAs, yielding a mixture of 5’monophosphates, and ^7me^GpppG-capped RNAs that were not affected by the phosphatase or PNK treatments. The other half of the phosphatase-treated RNA is incubated with a decapping enzyme, that will cleave-off the ^7me^Gppp-leaving a 5’terminal monophosphate. Both reactions are then subjected to an RNA ligation step using a 5′-adaptor RNA oligonucleotide linker. These two treatments resulted in the specific modification of the original noncapped or ^7me^GpppG capped viral RNAs with a 5′ adaptor linker (Fig.1A). The modified RNAs were then used as templates for the synthesis of cDNA via RT, using a reverse primer that hybridizes specifically to the positive strand of the viral RNA to be analyzed; afterwards, the RT reaction is used for qPCR reactions using oligos complementary to the 5’ adaptor linker sequence and to the 5’end of the viral RNA. For our purpose, specific oligos to detect an internal amplicon on the +RNA viral gene 10, and oligos for the 5’ adaptor linker sequence and the 5’end of the same viral RNA were used (Fig. 1B). This assay allows the quantification of the amount of ^7me^GpppG capped and noncapped (cap/noncap) viral RNAs in a given preparation.

To determine the presence or absence of the 5’cap structure on the positive strand of the genomic dsRNA, TLPs obtained from rotavirus infected MA104 cells were purified by isopycnic CsCl gradients. Before RNA extraction, the purified viral particles were treated with RNase A to remove any possible RNA bound to the exterior of the TLPs. Then, the genomic dsRNA was isolated and used to perform the enzymatic method for ^7me^GpppGcap quantification. We found that contrary to what was thought, there was a significant amount of noncapped RNAs present in the purified particles (40% +/- 8.3%, Fig. 1C-TLPs).

### The *in vitro* capping activity of VP3

Since there was a proportion of the encapsidated genomic dsRNA positive strands that was not capped, we evaluated the efficiency of the capping enzyme *in vitro*. After rotavirus entry into cells, the outermost layer of the TLPs is released, generating double layered particles (DLPs that are transcriptionally active. Under appropriate conditions purified DLPs can synthetize high amounts of viral +RNA *in vitro* (10), however, whether all mRNAs synthesized are capped is unknown. To characterize the efficiency of the VP3 capping activity, DLPs were obtained from RRV infected MA104 cells, the viral particles were concentrated by centrifugation through a 40% sucrose cushion and were resuspended in buffer TNE, which contains EDTA and causes the release of the outer layer proteins VP4 and VP7. These DLPs were used for an *in vitro* transcription assay as described in Materials and Methods; after ssRNA precipitation with LiCl, the proportion of ^7me^GpppGcap and noncapped 5’ends in the *in vitro* transcribed RNAs was determined as described above. A ratio of cap/noncap of 1:1 (50% +/- 9.1%, Fig. 1C) was found, suggesting that in this assay, VP3 has a capping efficiency of about 50%.

### VP3 capping activity during the infection cycle

To determine the capping activity of VP3 during the replication cycle, we quantitated the amount of ^7me^GpppG capped and noncapped viral RNA produced in infected cells. For this, total RNA was isolated from RRV-infected cells at different times post-infection, and the enzymatic assay to quantitate cap/noncap viral RNAs was performed. As shown in figure 2A, we found that the relative amount of capped versus noncapped viral +RNAs varied during the rotavirus replication cycle. The overall ratio of cap/noncap viral RNAs at the beginning of the infection (0 hpi) was 1:1 (50% +/- 1.8%), probably reflecting the cap/noncap ratio of the input virus, while at 3 hpi the RNA cap/noncap ratio decreased to 33% +/- 4.2%. In contrast, at 6 hpi the amount of ^7me^GpppG capped viral RNAs increased up to 73% +/- 4.2 %, and a ratio of cap/noncap RNAs close to 50% +/- 4.6% was observed at later times post-infection (9 and 12 hpi). In parallel, we quantitated by an RT-qPCR assay the relative abundance of viral +RNA gene 10 at the same times post-infection. In agreement with our previous findings (9), the amount of viral +RNA accumulated over time with a clear increase beginning at 6 hpi (Fig. 2B), suggesting that the ratio of cap/noncap RNA detected after 6 hpi represents mostly newly synthesized viral RNA. Together, these results suggest that the overall ratio of cap/noncap RNA is dynamic, varying during the early times post infection, while at 9 and 12 hpi it stabilizes to a 1:1 proportion, in agreement with the finding that purified virions and *in vitro* transcribed +RNAs also have a 1:1 proportion between capped and noncapped RNA molecules.

**Figure 2.**
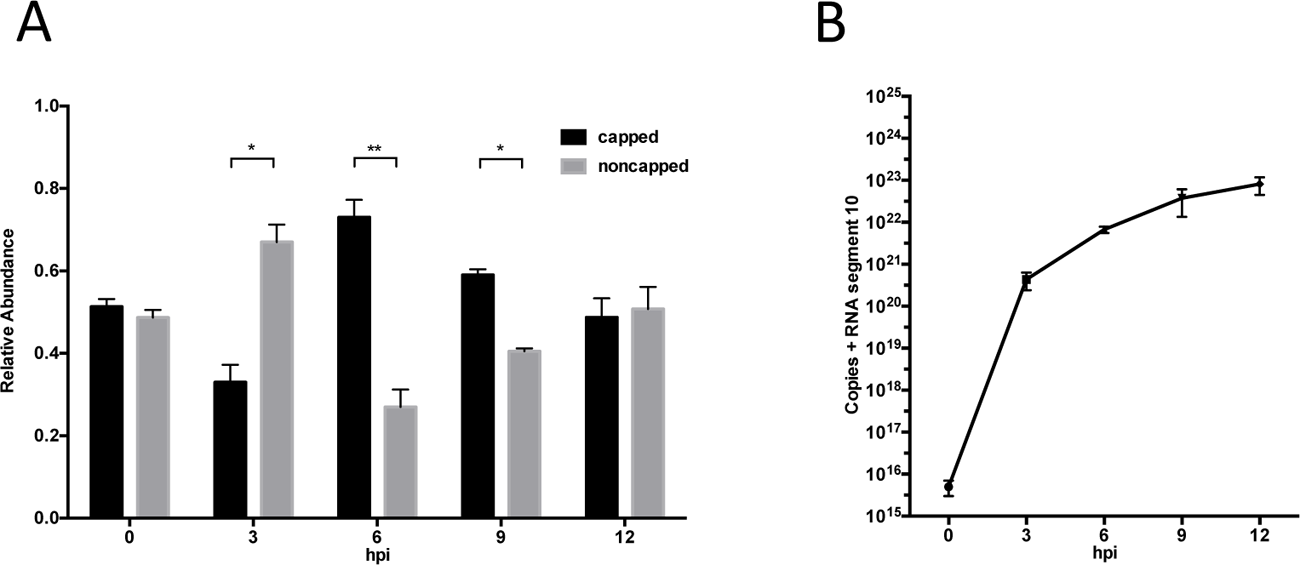
Quantitative determination of capped and noncapped RNAs at different times post-infection. A) MA104 cells were infected with RRV at an MOI of 3, total RNA was extracted at the indicated times post-infection, and the amount of capped and noncapped RNA was measured at each time point as indicated under Materials and Methods. The arithmetic means +/- standard deviation from three independent biological replicates are shown. B) The level of viral +RNA gene 10 synthesized in rotavirus infected cells were quantitated by an RT-qPCR assay at the indicated times post-infection. The number of viral copies at each time point was determined using a standard curve that was generated using a 10-fold serial dilution of an *in vitro* T7 transcribed RNA that contains the sequence of rotavirus RRV segment 10. The arithmetic means of +/- standard deviations from three independent biological replicates are shown. Statistical significance was determined by Student’s t test. *, P<0.05. **, P<0.01.

### Virus produced in the absence of VP3 is more infectious

We have previously reported that silencing the expression of VP3 through RNAi resulted in a decreased yield of infectious progeny, and also in a 10-fold reduction in the levels of mRNA and dsRNA during the infection, while paradoxically the synthesis of other viral proteins (besides VP3) was not affected (9). To evaluate the effect of the absence of VP3 on the encapsidation of viral RNAs (with and without cap), we purified viral particles produced in infected cells in which VP3 was silenced by RNAi, and determined their infectivity and the proportion of cap/noncap RNA present in the viral particles. For this, MA104 cells grown in 6-well plates were transfected either with an siRNA to VP3 or with an irrelevant siRNA for 72 h, and the cells were then infected with rotavirus RRV at an MOI of 3, and harvested at 12 hpi. The viral particles present in the cell lysates were concentrated by sedimentation through a 40% sucrose cushion as indicated under Materials and Methods, and resuspended in TNC buffer.

The efficiency of VP3 knock-down was analyzed by immunoblot analysis of total infected cell lysates and of semi-purified viral particles using a MAb directed to VP3, or a polyclonal antibody directed to TLPs (Fig. 3A). We found that while some VP3 could be detected in total cell lysates obtained from VP3-silenced cells, semi-purified particles contained very little, if any, detectable VP3. The same nitrocellulose blot was then stained with an antibody to TLPs, showing that there was a similar amount of viral proteins in the semi-purified particles (Fig. 3A). The yield of infectious virus produced under these conditions was determined by an immunoperoxidase focus-forming unit (FFU) assay (Fig. 3B), where we found that there was about a four-fold decrease in viral infectivity in the lysate of cells where VP3 was silenced compared to the lysate from the control condition. We also determined the specific infectivity of the semi-purified viral particles isolated under these conditions. Fig. 3C shows the results of these assays in which the viral titer of the viral particles obtained from cells in which VP3 was silenced or not, was expressed as a function of the concentration of protein present in the semi-purified particles from each condition. We found that the viral particles in which VP3 was silenced were about 46.5% +/- 14% more infectious relative to the non-silenced viral particles.

**Figure 3.**
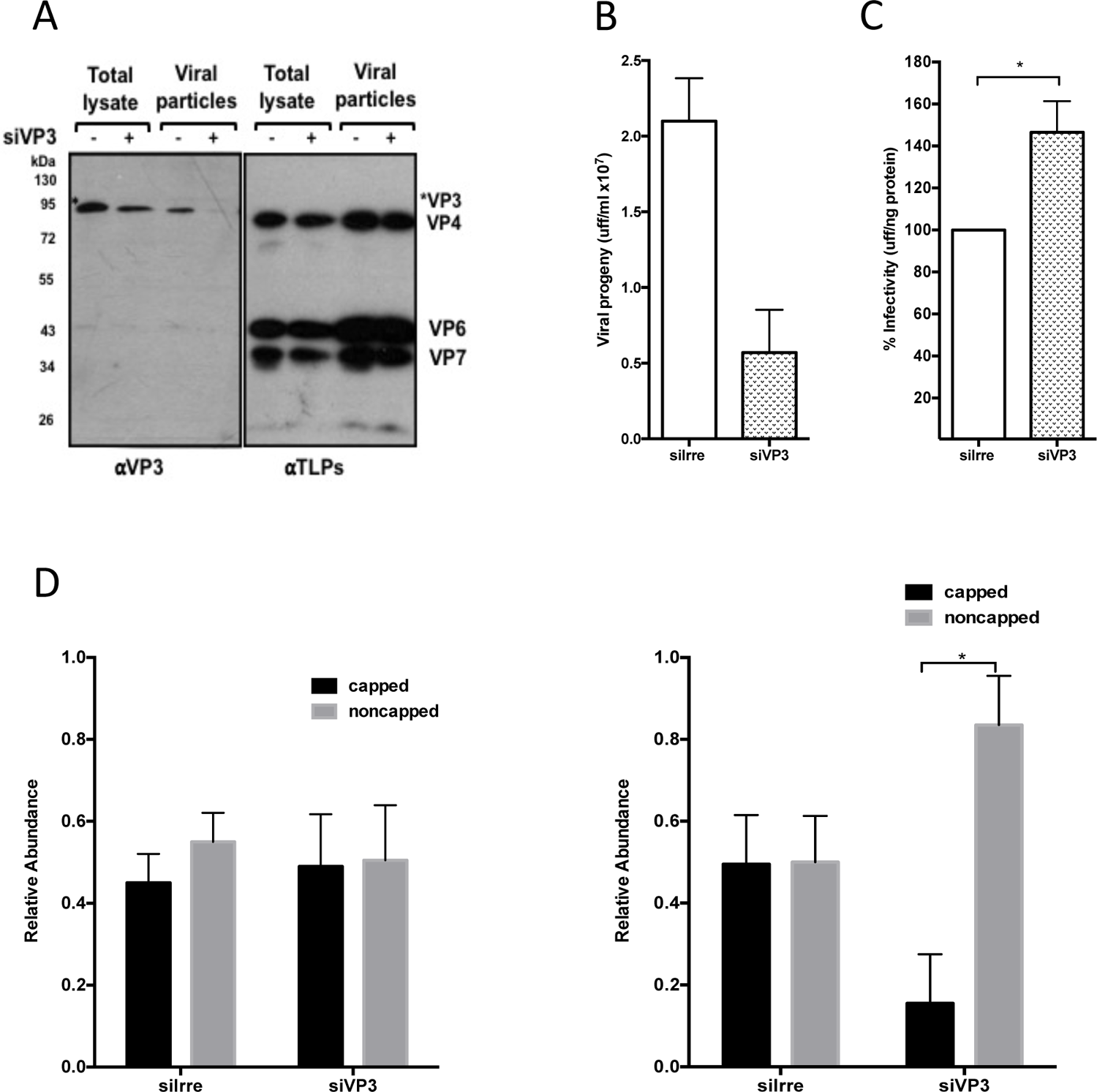
Virus produced in the absence of VP3 is more infectious. MA104 cells were transfected with the indicated siRNAs and 72 h post-transfection were infected with rotavirus RRV at an MOI of 3 and harvested at 12 hpi. The viral particles present in the cell lysates were concentrated by sedimentation through a 40% sucrose cushion and resuspended in TNC buffer. A) A representative immunoblot analysis of total cell lysates or viral particles from cells transfected with the indicated siRNAs. The expression of VP3, and the structural viral proteins VP4, VP6 and VP7 were detected with an anti-VP3 antibody (*α*VP3) or with an anti-TLP antibody (*α*TLP), respectively. **B)** The virus titer present in total cell lysates was determined by an immunoperoxidase focus-forming assay as described under Materials and Methods. Data are expressed as viral infectivity (uff/ml). The arithmetic means of +/- standard deviations from three independent experiments are shown. **C)** The viral titer in the semi-purified viral particles was determined by an immunoperoxidase focus-forming assay as described under Materials and Methods. The protein concentration in the viral particles obtained from each condition was determined as described under materials and methods. Data are expressed as the percentage of infectivity (uff/ng protein) obtained when the cells were transfected with an irrelevant siRNA (Irre) which was taken as 100% infectivity. The arithmetic means of +/- standard deviations from three independent experiments are shown. **D)** The relative amounts of capped and noncapped RNAs from the semi-purified particles treated with the indicated siRNA was determined as indicated under materials and methods. **E)** Quantitative determination of capped and noncapped of *in vitro* transcribed RNAs obtained from DLPs produced in infected cells treated with the indicated siRNA. Values shown represents means of results from three independent biological replicates with error bars representing standard deviation of the mean. Statistical significance was determined by Student’s t test. *, P<0.05.

To evaluate if the presence of ^7me^GpppGcap on the RNA correlated with the increased specific infectivity observed, the cap/noncap RNA ratio in the viral particles after VP3 knockdown was quantitated. For this, genomic dsRNA was purified from particles treated with either siVP3 or with siIrre, and was used for a capping determination assay. Prior to RNA extraction, viral particles were treated with RNase A to eliminate any possible viral +RNA contamination. Unexpectedly, we found that the ratio of cap/noncap RNA in the particles that were produced in VP3-knockdown cells was 1:1 (50% +/- 12.7%), a similar ratio to that found in the RNA extracted from particles obtained from the control condition (Fig. 3D). To corroborate that the semi-purified viral particles used in these assays lacked VP3 we explored the capping activity of the same viral particles; using an *in vitro* transcription assay that was performed as previously described. After RNA precipitation, the *in vitro* transcribed RNA was used for a capping determination assay. We found that in contrast to the 1:1 proportion of cap/noncap RNA obtained from particles treated with an irrelevant siRNA, only 15.5% +/- 12% of the RNA obtained from viral particles where VP3 was silenced had ^7me^GppGcap, while 83.5% +/- 12% of the in vitro synthesized RNA was not capped (Fig. 3E), indicating that these particles contained very little, if any VP3.

Together, these results show that in cells where the expression of VP3 was silenced, the yield of infectious viral particles is decreased compared to the control cells transfected with an irrelevant siRNA (Fig. 3B). Nevertheless, these particles have a higher specific infectivity (Fig. 3C), which does not seem to be related to the presence or absence of capped RNA since the genomic RNA enclosed in these particles had a similar cap/noncap RNA ratio (1:1). However, when these particles were used for an in *in vitro* transcription assay a higher proportion of noncapped RNAs was detected, underlining the absence of VP3. These results suggest that during genome encapsidation, RNAs with or without ^7me^GpppGcap are encapsidated in a similar proportion, irrespectively of the amount of capped RNAs present in infected cells.

### Effect of silencing the expression of RNase L and VP3 in viral RNA capping

It has been previously demonstrated that in addition to its capping activity, VP3 has a phosphodiesterase (PDE) activity that is involved in the degradation of 2’-5’oligoadenylates, thus preventing the activation of RNase L, an endoribonuclease that upon activation establishes a cellular antiviral state, cleaving single stranded regions of cellular and viral RNAs (24, 25). We previously showed that when the expression of VP3 and RNase L were co-silenced by RNAi in rotavirus-infected cells, the yield of infectious virus was higher and the integrity of the viral RNA increased compared to control conditions where irrelevant siRNAs were used (13). These results suggest that under conditions where the RNase L is silenced (and thus the PDE activity of VP3 was not needed), the capping activity of VP3 seems not to be essential for the production of infectious progeny virus. To analyze if the co-silencing of VP3 and RNase L influenced the abundance of capped viral RNA, MA104 cells grown in 48 well-plates and co-transfected with the indicated siRNAs, were infected with rotavirus strain RRV, and the monolayers were recovered at different times post-infection. The efficiency of co-silencing of VP3 and RNase L was evaluated by western blot using specific monoclonal antibodies against these proteins (Fig. 4A), and the infectivity of the viral particles produced under these conditions was determined at 12 hpi (Fig. 4B). As previously reported (13), we found that co-silencing the expression of VP3 and RNase L resulted in an increased viral infectivity of about 50%, which contrasts with the effect of only silencing VP3, conditions in which a lower total infectious titer was found compared with the virus titer obtained in control infected cells transfected with an irrelevant siRNA (Fig. 4B).

**Figure 4.**
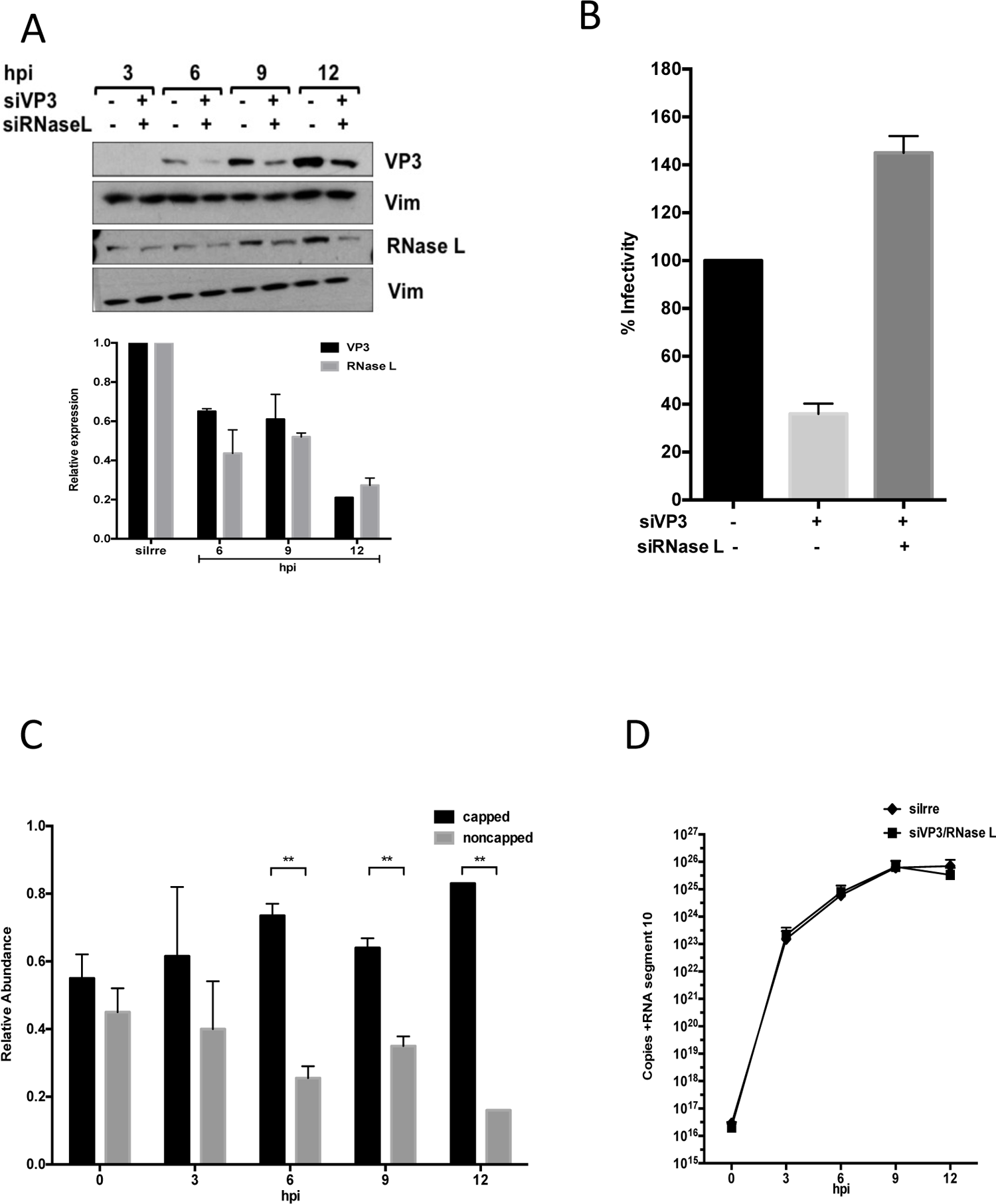
Effect of silencing the expression of RNase L and VP3 in viral RNA capping. MA104 cells were transfected with the indicated siRNA as indicated in materials and methods section. At 72 h post-transfection cells were infected with RRV at an MOI of 3 and were lysed at the indicated times post infection. **A)** A representative immunoblot analysis of total cell lysates from cells transfected with the indicated siRNAs. The expression of VP3, and RNase L were detected with the indicated antibodies, and anti-vimentin (Vim) was used as a loading control. Below the immunoblot a quantitation of the amount of protein detected under each condition is shown, the relative amount of the indicated proteins was calculated by densitometry of the bands using ImageJ software (LOCI, University of Wisconsin). Values represent the amount of protein detected as a percentage of the amount of protein detected in control cells treated with the irrelevant siRNA (siIrre) from three independent experiments. **B)** In parallel, cells transfected with the indicated siRNA were infected at an MOI of 3, at 12 hpi cells were lysed and the virus titer produced under these conditions was determined by an immunoperoxidase focus-forming assay as described. Data are expressed as the percentage of the infectivity obtained when the cells were transfected with an siIrre, which was taken as 100% infectivity. The arithmetic means +/- standard deviation of three independent experiments are shown. **C)** MA104 cells transfected and infected as in A) were harvested at the indicated times post-infection, total RNA was extracted, and the amount of capped and noncapped RNA was measured as indicated under Materials and Methods. The arithmetic means of +/- standard deviations from three independent experiments are shown. **D)** The levels of viral +RNA gene 10 synthesized in cells where VP3 and RNase L were co-silenced or in control cells transfected with an siIrre, were quantitated by an RT-qPCR assay at the indicated times post-infection. The number of viral copies at each time point was determined using a standard curve that was generated using a 10-fold serial dilution of an *in vitro* T7 transcribed RNA that contains the sequence of rotavirus RRV segment 10. The arithmetic means of +/- standard deviations from three independent biological replicates are shown. Statistical significance was determined by Student’s t test. *, P<0.05. **, P<0.01.

To quantitate the relative abundance of ^7me^GpppGcap and noncapped viral RNA produced under these conditions, total RNA was purified at different times post infection and analyzed in the capping assay as previously described. We found that at early times post-infection (0 and 3 hpi) the ratio of cap/noncap RNA was close to 1:1, which was also observed in control cells treated with an irrelevant siRNA (not shown). In contrast, starting at 6 hpi a higher proportion of capped RNA to noncapped RNA was found (73 +/- 3% at 6 hpi, 64 +/- 2% at 9 hpi and 83% +/- 2% at 12 hpi) (Fig. 4C). To demonstrate that there was not a decreased synthesis of positive-sense viral RNA under these conditions, the amount of viral RNA produced at the different time points of the assay was measured by RT-qPCR using specific oligos to detect viral gene 10. We found that at all time points measured a similar amount of segment 10 was produced under experimental and control conditions (Fig. 4D), suggesting that the differences found in the relative proportion of cap/noncap RNA in VP3-RNase L co-silenced infected cells were not due to differences in RNA transcription.

### The proportion of capped and non-capped RNA that is encapsidated in viral particles does not depend on the proportion present in infected cells

Since we found that the proportion of capped RNA produced in infected cells where VP3 and RNase L were co-silenced was higher than under control conditions, we investigated if this proportion was maintained in the newly formed viral particles. For this, viral particles produced in infected cells in which VP3 and RNase L were co-silenced were obtained as previously described (see Fig. 3); in these assays, as a control, viral particles were also obtained from cells in which only RNase L was knocked-down. The efficiency of silencing VP3 was analyzed by immunoblot analysis of the semi-purified viral particles using a MAb to VP3, or a polyclonal antibody directed to TLPs, and total infected lysates prior to particle isolation were used to demonstrate the silencing of RNase L (Fig. 5A).

**Figure 5.**
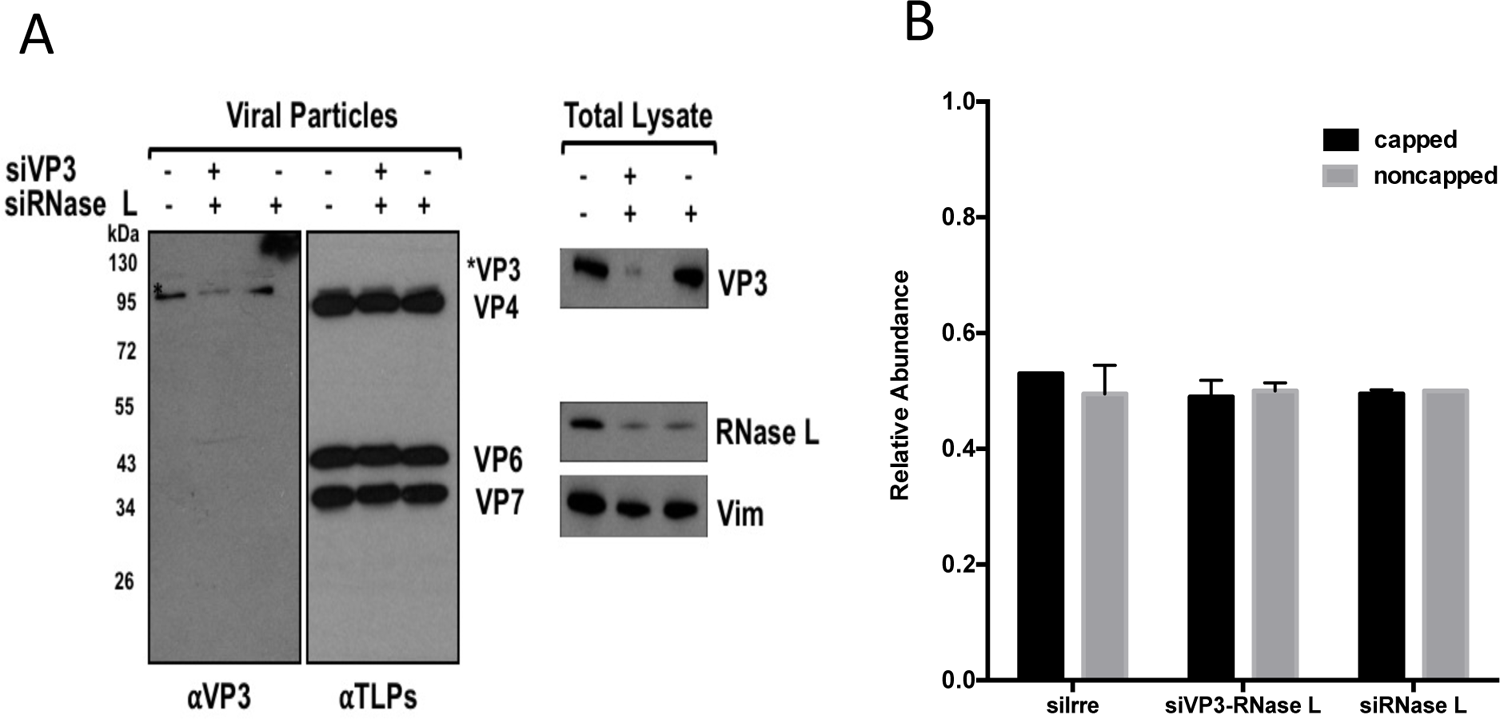
The proportion of capped and noncapped RNA that is encapsidated in viral particles does not depend on the proportion present in infected cells. MA104 cells were transfected with the indicated siRNA as indicated in Materials and Methods. At 72 h post-transfection cells were infected with rotavirus RRV at an MOI of 3 and harvested at 12 hpi. The viral particles present in the cell lysates were concentrated by sedimentation through a 40% sucrose cushion and resuspended in TNC buffer. **A)** The proteins were resolved by 10% of SDS-PAGE and detected by immunoblot analysis. A representative immunoblot of viral particles or total cell lysates from cells transfected with the indicated siRNAs is shown. The expression of VP3, and the structural viral proteins VP4, VP6 and VP7 in viral particles was detected with the indicated antibodies, respectively; and the expression of VP3, RNase L, and vimentin (Vim) in total cell lysates was detected with the indicated antibodies. **B)** The relative amounts of capped and noncapped RNAs from the semi-purified particles treated with the indicated siRNA was determined as indicated under Materials and Methods. The arithmetic means of +/- standard deviations from three independent biological replicates are shown.

The ratio of cap/noncap genomic positive-sense RNAs present in the viral particles obtained under these co-silencing conditions was quantitated. For this, the genomic dsRNA present in the semi-purified particles was extracted and used for a capping determination assay, as described. Prior to RNA extraction, viral particles were treated with RNase A to eliminate any possible viral +RNA contamination. We found that, contrary to what was expected, the ratio of capped to noncapped RNA in the particles that were produced in VP3/ RNase L co-silenced cells was 1:1 (50% +/- 2.0%), a similar ratio to that found in the RNA extracted from particles obtained from the control conditions in which an siRNA to RNase L or an irrelevant siRNA were used (Fig. 5B), suggesting that the proportion of capped and noncapped viral RNA that is encapsidated does not depend on the relative abundance of these RNA populations in the infected cells.

## DISCUSSION

In the celĺs cytoplasm, noncapped RNAs are detected by RIG I and MDA5, proteins of the innate immune system (RIG I and MDA5), that recognize specific epitopes present on the 5’end of non-self RNAs, including the presence of tri-, di- and monophosphate groups. The recognition of these epitopes triggers a signaling cascade that leads to the expression of interferon (IFN) as an initial immune response against viral infection (26, 27). Viruses have evolved different mechanisms to overcome this recognition, thus preventing the induction of IFN (28). Rotaviruses have different strategies to avoid the recognition of the dsRNA genome produced during the replication cycle; however, even though the viral capsid contains a protein (VP3) with all the enzymatic activities required to add the cap structure to the 5’ end of the viral transcripts, here we have found that during the virus replication cycle noncapped RNAs are produced, and the molecular sensors RIG-I and MDA-5 are activated (29)(30); the continuous production of noncapped RNAs throughout the viral cycle might explain, at least in part, the activation of these cellular sensors.

In this work, we quantitated the presence of ^7me^GpppGcap on rotavirus viral RNAs employing a modified enzymatic assay previously used in the characterization of the 5’ ends of Sindbis virus. Counterintuitively, we found a significant proportion of noncapped +RNAs present in the mature infectious particles (TLPs); interestingly, in *in vitro* assays, a similar proportion (1:1) of 5’capped and noncapped RNAs was synthesized, suggesting that the efficiency of the capping enzyme is not 100%, as was previously observed by Uzri et al., who found that in viral RNA isolated from rotavirus-infected cells and in *in vitro* transcribed RNA there was a significative proportion of RNA molecules with exposed 5’ phosphate groups and incompletely 2’-O-methylated cap structures (31). In their study, the absence of cap was determined by an indirect approach in which the viral RNAs obtained either from *in vitro* transcription or isolated from infected cells were transfected into murine embryonic fibroblasts (MEFs), and the level of secreted IFN-beta was measured by ELISA (31). In this work we found that the abundance of ^7me^GpppGcap and noncapped RNAs in rotavirus-infected cells was dynamic since their proportions varied at different times post infection, with a higher amount of capped RNAs at 6 and 9 hpi. The increased proportion of capped RNAs found at these times could be the result of the increased transcription observed during the replication cycle (Fig. 2B) (9); while at 12 hpi the proportion of capped and noncapped RNAs was close to 1:1, which was the same proportion of ^7me^GpppGcap to noncapped RNA detected in the genomic +RNA present in the mature viral particles (Fig. 1C).

Interestingly, besides its capping activity VP3 has several mechanisms to counteract the innate immune response; it has been recently shown that VP3 is involved in the degradation of MAVS, blocking the IFN-*λ* production in rotavirus-infected intestinal epithelial cells, thus evading the host cell immune response (32). It also has a phosphodiesterase (PDE) domain that degrades the 2’-5’oligoadenylates produced in response to the detection of dsRNA by the OAS/RNase L pathway; this PDE activity prevents the activation of RNase L, an enzyme that is in charge of degrading cellular and viral RNAs in infected cells to restrain viral infection (24). Thus, apparently VP3 is a multifunctional protein that can counteract the activation caused by the lack of cap in the transcripts synthesized during the infection.

The effect of silencing the expression of VP3 on the synthesis of viral proteins and on the production of viral particles has been reported; however, how the addition of cap on viral RNAs affects these processes had not been characterized (9, 13). We found that even though under conditions where VP3 is silenced there is a significant lower amount of RNA, the translation of viral proteins is not affected, and the viral particles produced (although in lesser amounts) contain a similar ratio of capped and noncapped RNAs compared to those obtained from cells treated with an irrelevant siRNA. Most probably, all the ^7me^GpppGcap containing viral RNAs produced in the VP3 silenced infected-cells originate from the incoming particles, and might be enough to direct the synthesis of the viral proteins during the infection cycle, and to be encapsidated in newly formed viral particles (although in lesser amount). On the other hand, the particles produced under these conditions, although less abundant, had a slightly higher specific infectivity than those obtained in control cells. It is tempting to suggest that these particles probably have a better sorting of capped/noncapped RNA segments, as opposed to control infected cells where probably there could be more viral particles assembled containing on average a higher proportion of uncapped RNAs resulting in a relative lower specific infectivity. As expected, the viral particles assembled in VP3 silenced cells contained very little, or no VP3, since the *in vitro*-transcribed RNA from DLPs produced under these conditions had a significantly lower amount of ^7me^GpppGcap RNA. It is interesting to note that even though there is little or no VP3 in these particles, viral RNA was encapsidated, suggesting that VP3 might not play a role in this function, contrary to what has been proposed.

We have previously found that in cells where the expression of VP3 and RNase L were co-silenced, the yield of viral infectious particles was similar to that obtained from control cells where no genes were silenced, suggesting that the capping activity of VP3 was not essential for the production of infectious progeny virus under these conditions (13). Here we found that there is a higher proportion of capped to uncapped viral RNAs produced in the infected cells where VP3 and RNase L were co-silenced, such that by 12 hpi, more that 80% of the viral +RNA is capped, and yet the proportion of ^7me^GpppGcap and noncapped +RNA found in the viral particles isolated from these conditions are in a 1:1 proportion, suggesting that there should be a mechanism that controls the proportion of capped:uncapped +RNA that is encapsidated, despite the levels of ^7me^GpppGcap RNA present in rotavirus-infected cells.

In contrast, in reovirus, another member of the family *Reoviridae*, it has recently been shown that the efficient capping of viral RNAs, in addition to its role in enhancing protein translation, has a function in promoting the assembly of particles and the genomic RNA incorporation into progeny virions (33). These results were obtained in a reverse genetic assay of reovirus, in which the role of C3P3 a fusion protein containing the T7-RNA polymerase and the African swine fever virus NP868R capping enzyme was tested for its ability to rescue reovirus; it remains to be determined if the activity of the native capping reovirus enzymes during the replication cycle of this viruses behaves similarly.

Recently, it was reported that noncapped RNAs have an important relevance during SINV infection in an *in vitro* and *in vivo* model, since when the viral protein involved in cap addition (nsP1) was mutated to decrease the amount of cap on viral RNAs a slight decrease in mortality of infected mice was observed. Conversely, an increased addition of cap resulted in an almost complete abrogation of mortality and morbidity compared to mice infected with a wild type virus (34, 35). Thus, in SINV noncapped RNAs play a critical role in determining whether the virus is neurovirulent or not, however if the cap structure has a role in genome encapsidation is not known. Whether there is a role of the non-capped viral rotavirus RNAs early during the infection needs to be determined.

Since the domains involved in cap addition of VP3 have been recently defined (11, 12), and taking advantage of the available reverse genetic system for rotavirus (36), it would be interesting to generate VP3 mutants that impair some of the regions involved in the different functions of the methyl-guanylyl transferase activity of this protein and establish if ^7me^GpppGcap and noncapped RNAs have a role during rotavirus infection and if the capping activity of VP3 is important for particle assembly.

## ACKNOWLEDGMENTS

We are grateful to J.L. Reyes for providing the 5’RNA adapter, to R. Espinosa and M.A. Espinoza for their excellent technical assistance, and to K.J. Sokolosky for his help with the analysis of the results and to C. Sandoval-Jaime for his valuable input during the development of this project. The work of P. Gaytán, E. López, and J. Yañez from the DNA sequencing and synthesis core unit is also acknowledged. This work was supported by grants A1-S-15356 and 302965 from CONACyT. JMC was a recipient of a scholarship from CONACyT

## REFERENCES

1. Ramanathan A, Robb GB, Chan S-H. 2016. mRNA capping: biological functions and applications. Nucleic Acids Res2016/06/17. 44:7511–7526.

2. Galloway A, Cowling VH. 2019. mRNA cap regulation in mammalian cell function and fate. Biochim Biophys acta Gene Regul Mech 2018/10/09. 1862:270–279.

3. Furuichi Y. 2015. Discovery of m(7)G-cap in eukaryotic mRNAs. Proc Jpn Acad Ser B Phys Biol Sci 91:394–409.

4. Decroly E, Ferron F, Lescar J, Canard B. 2012. Conventional and unconventional mechanisms for capping viral mRNA. Nat Rev Microbiol 10:51–65.

5. Jan E, Mohr I, Walsh D. 2016. A Cap-to-Tail Guide to mRNA Translation Strategies in Virus-Infected Cells. Annu Rev Virol 3:283–307.

6. Troeger C, Khalil IA, Rao PC, Cao S, Blacker BF, Ahmed T, Armah G, Bines JE, Brewer TG, Colombara D V., Kang G, Kirkpatrick BD, Kirkwood CD, Mwenda JM, Parashar UD, Petri WA, Riddle MS, Steele AD, Thompson RL, Walson JL, Sanders JW, Mokdad AH, Murray CJL, Hay SI, Reiner RC. 2018. Rotavirus Vaccination and the Global Burden of Rotavirus Diarrhea among Children Younger Than 5 Years. JAMA Pediatr 172:958–965.

7. Estes M.K KA. 2007. Rotaviruses, p. 1917–1974. In Wilkins, WKHW& (ed.), Fields Virology, 5th ed. In Knipe DM, Howley PM, Griffin DE, Lamb RA, Martin MA, Roizman B, Straus SE (ed), Philadelphia.

8. Patton JT, Spencer E. 2000. Genome Replication and Packaging of Segmented Double-Stranded RNA Viruses. Virology 277:217–225.

9. Ayala-Breton C, Arias M, Espinosa R, Romero P, Arias CF, López S. 2009. Analysis of the kinetics of transcription and replication of the rotavirus genome by RNA interference. J Virol2009/06/24. 83:8819–8831.

10. Chen D, Luongo CL, Nibert ML, Patton JT. 1999. Rotavirus Open Cores Catalyze 5′-Capping and Methylation of Exogenous RNA: Evidence That VP3 Is a Methyltransferase. Virology 265:120–130.

11. Kumar D, Yu X, Crawford SE, Moreno R, Jakana J, Sankaran B, Anish R, Kaundal S, Hu L, Estes MK, Wang Z, Prasad BVV. 2020. 2.7 Å cryo-EM structure of rotavirus core protein VP3, a unique capping machine with a helicase activity. Sci Adv 6:eaay6410.

12. Ogden KM, Snyder MJ, Dennis AF, Patton JT. 2014. Predicted structure and domain organization of rotavirus capping enzyme and innate immune antagonist VP3. J Virol2014/06/04. 88:9072–9085.

13. Sánchez-Tacuba L, Rojas M, Arias CF, López S. 2015. Rotavirus Controls Activation of the 2’-5’-Oligoadenylate Synthetase/RNase L Pathway Using at Least Two Distinct Mechanisms. J Virol2015/09/23. 89:12145–12153.

14. Zhang R, Jha BK, Ogden KM, Dong B, Zhao L, Elliott R, Patton JT, Silverman RH, Weiss SR. 2013. Homologous 2’,5’-phosphodiesterases from disparate RNA viruses antagonize antiviral innate immunity. Proc Natl Acad Sci U S A 110:13114–9.

15. Sokoloski KJ, Haist KC, Morrison TE, Mukhopadhyay S, Hardy RW. 2015. Noncapped Alphavirus Genomic RNAs and Their Role during Infection. J Virol 2015/04/01. 89:6080–6092.

16. Pando V, Iša P, Arias CF, López S. 2002. Influence of calcium on the early steps of rotavirus infection. Virology 295:190–200.

17. Montero H, Arias CF, Lopez S. 2006. Rotavirus Nonstructural Protein NSP3 is not required for viral protein synthesis. J Virol 80:9031–8.

18. López T, Camacho M, Zayas M, Nájera R, Sánchez R, Arias CF, López S. 2005. Silencing the morphogenesis of rotavirus. J Virol 79:184–192.

19. Martínez JL, Arnoldi F, Schraner EM, Eichwald C, Silva-Ayala D, Lee E, Sztul E, Burrone ÓR, López S, Arias CF. 2019. The Guanine Nucleotide Exchange Factor GBF1 Participates in Rotavirus Replication. J Virol 93:e01062–19.

20. Gutiérrez M, Isa P, Sánchez-San Martin C, Pérez-Vargas J, Espinosa R, Arias CF, López S. 2010. Different rotavirus strains enter MA104 cells through different endocytic pathways: the role of clathrin-mediated endocytosis. J Virol2010/07/14. 84:9161–9169.

21. Livak KJ, Schmittgen TD. 2001. Analysis of Relative Gene Expression Data Using Real-Time Quantitative PCR and the 2−ΔΔCT Method. Methods 25:402–408.

22. Jenni S, Salgado EN, Herrmann T, Li Z, Grant T, Grigorieff N, Trapani S, Estrozi LF, Harrison SC. 2019. In situ Structure of Rotavirus VP1 RNA-Dependent RNA Polymerase. J Mol Biol2019/06/21. 431:3124–3138.

23. Hauser M, Dearnaley WJ, Varano AC, Casasanta M, McDonald SM, Kelly DF. 2019. Cryo-EM Reveals Architectural Diversity in Active Rotavirus Particles. Comput Struct Biotechnol J 17:1178–1183.

24. Zhou A, Hassel BA, Silverman RH. 1993. Expression cloning of 2-5A-dependent RNAase: A uniquely regulated mediator of interferon action. Cell 72:753–765.

25. Schwartz SL, Conn GL. 2019. RNA regulation of the antiviral protein 2’-5’-oligoadenylate synthetase. Wiley Interdiscip Rev RNA2019/04/15. 10:e1534–e1534.

26. Kell AM, Gale M. 2015. RIG-I in RNA virus recognition. Virology 479–480:110– 121.

27. Brisse M, Ly H. 2019. Comparative Structure and Function Analysis of the RIG-I-Like Receptors: RIG-I and MDA5. Front Immunol.

28. Beachboard DC, Horner SM. 2016. Innate immune evasion strategies of DNA and RNA viruses. Curr Opin Microbiol2016/06/08. 32:113–119.

29. Broquet AH, Hirata Y, McAllister CS, Kagnoff MF. 2011. RIG-I/MDA5/MAVS Are Required To Signal a Protective IFN Response in Rotavirus-Infected Intestinal Epithelium. J Immunol 186:1618 LP – 1626.

30. Sen A, Pruijssers AJ, Dermody TS, García-Sastre A, Greenberg HB. 2011. The Early Interferon Response to Rotavirus Is Regulated by PKR and Depends on MAVS/IPS-1, RIG-I, MDA-5, and IRF3. J Virol 85:3717–3732.

31. Uzri D, Greenberg HB. 2013. Characterization of Rotavirus RNAs That Activate Innate Immune Signaling through the RIG-I-Like Receptors. PLoS One 8:1–14.

32. Ding S, Zhu S, Ren L, Feng N, Song Y, Ge X, Li B, Flavell RA, Greenberg HB. 2018. Rotavirus VP3 targets MAVS for degradation to inhibit type III interferon expression in intestinal epithelial cells. Elife 7:e39494.

33. Eaton HE, Kobayashi T, Dermody TS, Johnston RN, Jais PH, Shmulevitz M, López S. 2017. African Swine Fever Virus NP868R Capping Enzyme Promotes Reovirus Rescue during Reverse Genetics by Promoting Reovirus Protein Expression, Virion Assembly, and RNA Incorporation into Infectious Virions. J Virol 91:e02416–16.

34. LaPointe AT, Landers VD, Westcott CE, Sokoloski KJ, Patton JT. 2020. Production of Noncapped Genomic RNAs Is Critical to Sindbis Virus Disease and Pathogenicity. MBio 11:e02675–20.

35. LaPointe AT, Moreno-Contreras J, Sokoloski KJ. 2018. Increasing the Capping Efficiency of the Sindbis Virus nsP1 Protein Negatively Affects Viral Infection. MBio 9:e02342–18.

36. Kanai Y, Komoto S, Kawagishi T, Nouda R, Nagasawa N, Onishi M, Matsuura Y, Taniguchi K, Kobayashi T. 2017. Entirely plasmid-based reverse genetics system for rotaviruses. Proc Natl Acad Sci 114:2349 LP – 2354.

